# Characterization and identification of enzyme-producing microflora isolated from the gut of freshwater Prawn, *Macrobrachium* Spp

**DOI:** 10.1101/2024.12.10.627853

**Authors:** Dhanraj N Nimbalkar, Virdhaval M. Nalavade, Jaykumar J. Chavan

## Abstract

Isolation of extracellular enzyme degrading aerobic bacteria in the digestive tracts of fresh water prawn was carried out. Gut bacteria were isolated on carboxymethyl cellulose agar plates, starch agar media plates, gelatine peptone agar media plates. The isolated strain was qualitatively screen based on their extracellular enzyme on selective media. The strain was characterised based on morphological, physiological and biochemical characterization identified Bacillus species. Bacillus species was isolated by colony characterization carried out by using Gelatin agar Media, carboxymethyl cellulose media and congo red CMC media and starch agar media for different enzyms. Isolates are capable of hydrolysing proteins and carbohydrates indicating their importance in fish nutrition.

## 1. INTRODUCTION

Gut microorganisms play an important role in the digestion and assimilation of food consumed by their host animals.

Prawn receives bacteria in the digestive tract from the aquatic environment through water and food that are populated with bacteria. Being rich in nutrient, the environment of the digestive tract of shrimp confers a favourable culture environment for the microorganisms. In the present study, an attempt has been made to investigate the relative amount of protease, amylase, cellulaseproducing bacteria in the gastrointestinal tract of prawn (*Fenneropenaeusindicus*). Further, intestinal isolates were evaluated for extracellular enzyme producing capacities.

Data suggests that the composition of the specific enzyme producing bacterial flora in the Prawn digestive tract may have correlation with their feeding habits. It is a consensus view that dense bacterial population levels occurs in the gastrointestinal tract of Prawn. The gut microflora may be categorized as either autochthonous (indigenous) or allochthonous (transient) depending upon its ability to colonize and adhere to the mucus layer in the digestive tract of Prawn. Prawn gut bacteria may be used as probiotics for Prawn. Those occurring in the gastrointestinal tract of Prawn. One of the major criteria for selecting a probiotic strain is its ability to adhere and colonize the digestive tract of the host.

The isolation and enumeration of heterotrophic bacteria from the gastrointestinal tract of fresh water Prawn *Fenneropenaeusindicus*and were carried out to determine their importance in host Prawn. Amyllytic, proteolytic and cellulolytic bacteria were detected in the Prawn gut. Among specific enzyme producing bacteria, proteolytic and cellulolytic bacteria were present in higher numbers within the gut of Prawn. Selected intestinal isolates were analysed for extra cellular enzyme production capacities. Protease and cellulose activities were exhibited by all the bacterial isolates. The result of this study indicated the probability of digestive enzyme supplementation by the intestinal bacteria in the Prawn gut. The information generated from the present investigation might contribute to the utilization of these extracellular enzyme producing bacteria in the commercial aquaculture. It seems logical that the gut bacteria have a role in nutrition, growth and disease susceptibility in Prawn as it has been established for homothermic species. And understanding of the indigenous microbiota in Prawn may help to improve feeding and other conditions for the intensive rearing of fish.The purpose of this research was to isolate and access the enzyme producing microbes from the prawn gut.

## 2. MATERIALS AND METHODS

### Study area and collection of water sample

The Vennariver is one of the important tributary of the river Krishna originated from Sahayadri mountain ranges in Mahabaleshwar, Satara District, Maharashtra. The sampling area was Venna river basin near Sataracity.

Water, sediment and adult *M. rosenbergii* samples were collected from above mentioned location (Figure 1) and necessary precautionswere taken to minimize the contamination of the sample. Water andsediment samples were collected in sterile bottles and sterile jarsrespectively. The adult *M. rosenbergii*were collected by fisherman inlive condition and brought to the laboratory for analysis. All the samples exceptthe adult *M. rosenbergii*were kept in an icebox and immediatelybrought to the laboratory for analysis.

### Analysis of the Physio-Chemical Parameters

Physico-chemical parameters of water samples such astemperature were measured *in situ* using centigrade thermometer,salinity by salinity refractometer (Atago, Japan) and pH by digitalpH meter (Eutech, Singapore). Dissolved oxygen was estimated bythe Winkler method [15]. The pH of sediment was measured usingthe method described by Sharmila et al. [16].Other parameters such as hardness, TDS, phosphate, nitrate, chloride, alkalinity, COD, BOD, DO etc. were estimated by using standard methods described by APHA (1985), Trivedy and Goel (1986), Kodarkar (2006).

### Isolation of enzyme producing bacteria from gut flora

The test samples of *M. rosenbergii*were dissected to remove intestine aseptically, weighed andhomogenized in glass homogeniserwith sterilized 0.85% NaCl solution (10:1; volume:weight)and serially diluted to 10^-6^.To isolate the hetrophic bacterial population, diluted samples (0.1 ml) were streaked aseptically on sterilized media plates & incubated over night at 28°C. The well separated colonies that apparently have different morphological appearance (e.g. colony shape, colour, elevation) were isolated & streaked separately on specific media plates to obtain pure culture.For amylase enzyme activity, media containing agar-agar with 1% starchwas prepared aseptically.The cellulase agar media was prepared with 1% carboxymethyl cellulose aseptically.The protease gelatin peptone agar mediumwas prepared with 1% gelatinaseptically.Enzyme producers were is visualized by translucentzone or clear zone around the colonies.

### Identification of bacteria

#### Morphological analysis, colony characteristics and motility

Morphological characterization of isolated contaminants was done on the basis of simple staining and gram staining methods (Sherman, 2005). Colonies of isolates were screened for characters like size, shape, colour, margin, elevation, opacity, consistency.To study the motility 48 hours old culture was taken for observing motility by hanging drop method under oil immersion objective. The contaminating fungi were identified by colony characteristics (appearance, colour and pigmentation), morphology of vegetative hyphae and macroconidia produced by comparing with standards enlisted by (Barnett and Hunter).

#### Biochemical analysis

Biochemical characterization of isolates was done by using a series of biochemical tests, like Nitrate reduction, Catalase, Starch hydrolysis, Casein hydrolysis, Gelatin hydrolysis, Hydrogen Sulphide gas (H_2_S) production, sugar fermentation ability with acid and gas production for various isolates (Sherman, 2005). The characterized bacterial strains were compared with standard in Bergey’s Manual of Bacteriology [Bergey,1994 and Buchanan 1984, Holt,1994].

#### PCR Amplification of the 16S rDNA, Sequencing and Phylogenetic Analysis of Probiotic Bacteria

DNA extraction was done by Bacterial Genomic DNA (prep) Kit(Chromous Biotech, India) and 16S rDNA genes were amplified withthe universal primers 27f (5’-AGAGTTTGATCCTGGCTCAG-3’) and1492r (5’-GGTTACCTTGTTACGACTT-3’) by using the polymerasechain reaction (PCR) [21].The nearest taxa of the 16S rRNA gene sequence (1418–1542bases) was identified by BLAST sequence similarity search (https://blast.ncbi.nlm.nih.gov/Blast.cgi). The CLUSTAL W software wasused to align 16S rRNA gene sequences and Maximum Likelihood (ML) and Neighbour-Joining (NJ) methods with MEGA version 5[22] were used to construct the phylogenetic tree.

#### Determination of antibacterial activity of isolates by welldiffusion method

For the determination ofantibacterial activity by well diffusion method, 2 ml of a youngculture (16–18 hour in TSB) of*V. harveyi*,*V. vulnificus, S. typhi*and *E. coli*, were preparedand poured over the TSA medium and incubated at 30°C for 15minutes. Sterile gel puncher was used to punch three millimeter diameter wells in the plates and 30 microliters of 18 hours bacterialculture in TSB was pipette into the wells and incubated at 30°C for24 hours. Clear zone around the wells was noted as the presence ofantibacterial activity.

#### In vivo experiment for the safety of probiotic strain topost larvae of *M. rosenbergii*

Testing the safety of probioticstrain used is important hence for this themixed culture of potential probioticbacteria were added to 1 L beaker containing 500 ml sterilized filtered freshwater to obtain 10^5^ cells ml^-1^. To each beaker, which 25numbers of *M. rosenbergii*post larvae were added andstudied for any signs of disease or mortality up for 96 hours period.The control set consist of 1 L beaker containing 500 ml sterilized filtered freshwaterwithout any bacterial inoculum.The experiments were performed in triplicate and observations were recorded.

## RESULT AND DISCUSSION

### Analysis of the Physio-Chemical Parameters

Physico-chemical parameters of water and sediment samplescollected from the sampling sites revealed that temperature, pHand DO of water ranged between 28.5°–31.0°C, 5.8–6.7 and 6.9–7.2mg L-1 respectively. The salinity of water samples were within therange of 0–4 ppt. The pH of sediment samples ranged between 5.48–6.46. Similar pH values were reported by Nandan and Unnithan[23] from Vembanad Lake and the physico-chemical parametersanalysed were within the optimal range for growth and survival of*M. rosenbergii*in their natural environment [14]. Low salinity andthe slightly acidic nature of water and sediment samples recordedin the present study are in agreement with the results [24] fromthe Kuttanad region of Vembanad Lake. Thanneermukam barrageconstructed across the Vembanad Lake to prevent the ingress ofsaline water into the rice fields of this area cause the low salinityof water.

| Sr.No. | Test | Observations |
| --- | --- | --- |
| 1 | pH | 7.5 |
| 2 | EC | 0.27 |
| 3 | Na<sup>+</sup> | 49.14 |
| 4 | K<sup>+</sup> | 0.78 |
| 5 | CO<sub>3</sub> | ----- |
| 6 | HCO<sub>3</sub> | ----- |
| 7 | Cl | 2.2 |
| 8 | TSS | 5000 |
| 9 | TDS | 8000 |
| 10 | Hardness | 22.4 |
| 11 | Alkalinity | 160 |
| 12 | DO | 12.4 |
| 13 | BOD | ----- |
| 14 | COD | ----- |
| 15 | Temperature | 29.7 |

### Identification of enzyme producing bacterial isolates

The isolated three bacteria were analysed for their morphology, colony characteristic, motility and various biochemical characters (Table 1 & 2) according to BERGY’S manual (Bergey,1994 and Buchanan 1984, Holt,1994).

**Table 1.** Morphology, colony characteristic and motility of isolated microbial contaminants.

| Particulars | Isolated Bacteria |  |  |
| --- | --- | --- | --- |
|  | Isolate 1 | Isolate 2 | Isolate 3 |
| **Colony Characteristics** |  |  |  |
| Size | 1.0 mm | 1.0 mm | 1.0 mm |
| Shape | Circular | Circular | Circular |
| Colour | Cream | Light cream | Cream |
| Margin | Entire | Entire | Entire |
| Elevation | Convex | Convex | Flat |
| Opacity | Opaque | Opaque | Opaque |
| Consistency | Moist | Smooth | Smooth |
| Gram staining | Positive (violet rods) | Negative (pink rods) | Negative (pink rods) |
| Motility | Motile | Motile | Motile |

**Table 2.** Various biochemical characters of isolated microbial contaminants.

| Particulars | Isolated Bacteria |  |  |
| --- | --- | --- | --- |
|  | Isolate 1 | Isolate 2 | Isolate 3 |
| **Sugar Fermentation Capability** |  |  |  |
| Glucose | + | - | - |
| Sucrose | + | - | + |
| Fructose | + | - | - |
| Lactose | + | + | - |
| Mannose | + | - | + |
| Dextrose | + | - | + |
| Trehalose | + | + | + |
| Xylose | + | - | - |
| Sorbitol | + | - | - |
| Arabinose | + | - | - |
| **Other biochemical characters** |  |  |  |
| Starch | + | + | + |
| Catalase | + | + | + |
| Gelatin | - | + | + |
| Nitrate Reduction | - | - | - |
| Production of H<sub>2</sub>S | - | - | - |
| Casein | - | + | + |

According to Morphological, physiological and biochemical characterization of amylase isolatescolony Characters shows tentative genus of isolates is

| Isolates | Genus | Enzyme activity |
| --- | --- | --- |
| Isolates A | *Bacillus* Spp. | Cellulase |
| Isolates B | *Pseudomonas* Spp. | Protease |
| Isolates C | *Pseudomonas aruginosa* | Amylase |

### Antibiotic sensitivity test

Various antibiotics like Cefotaxime, Gentamycin, Kanamycin and Tetracycline at following concentrations 50, 100, 150, 200 and 250 mg/L in LB medium were used to study the growth inhibition of isolated six microbial contaminants microbial growth. It was observed that addition of Cefotaxime at concentration of 250 mg/ L was satisfactory to control the growth of all six microbial contaminants (Table 4).

**Table 4:** Growth of isolated microbial contaminants on various antibiotic concentrations.

| Sr. No. | Test Bacteria | Bacterial Isolates |  |  |
| --- | --- | --- | --- | --- |
|  |  | *Bacillus* Spp. | *Pseudomonas* Spp. | *Pseudomonas aruginosa* |
| 1 | *V. harveyi* | ++ | ++ | + |
| 2 | *V. vulnificus* | + | + | ++ |
| 3 | *S. typhi* | ++ | + | + |
| 4 | *E. coli* | + | + | +++ |

**Note:** +++: No inhibition, ++: Less Inhibition, +: Maximum Inhibition

### In vivo experiment for the safety of probiotic strain topost larvae of *M. rosenbergii*

Before going for the experimental trial, it should be confirm thatthe probiotic bacteria should not show any pathogenic or adverseeffect on host [65] and the probiotic bacteria are then selectedaccording to the antagonistic property against the pathogens andby *in vitro* testing [66–70]. The results of pathogenicity effect of the*Brevibacilluslaterosporus*on post larvae of *M. rosenbergii*(Figure5) showed that the bacteria has no pathogenic effect on the postlarvae and the survival was higher with that of control (*p* < 0.01).

## Conclusion

In the study, an attempt was made to find out the bacterial diversity associated with various life stages of giant freshwater prawn, and screening for probiotic potential among these bacteria. The study strengthens the microbial ecology and correlation of microbial communities (microorganisms on water, sediment, gut and larvae). The screening and probiotic potential study found that *Brevibacillu slatrosporus* showed antibacterial activity against fish and prawn pathogens. No adverse effect was noticed when the PL of *M. rosenbergii* challenged with the selected probiotic strains. The present study suggested *Brevibacillus laterosporus* as a promising probiotic candidate for hatchery and culture operations of *M. rosenbergii*. However, thorough studies are suggested with detail evaluation on the effect in vivo with these bacteria.

